# A VPS13D spastic ataxia mutation disrupts the conserved adaptor binding site in yeast Vps13

**DOI:** 10.1101/768366

**Authors:** Samantha K. Dziurdzik, Björn D. M. Bean, Michael Davey, Elizabeth Conibear

## Abstract

Mutations in each of the four human VPS13 (VPS13A-D) proteins are associated with distinct neurological disorders: chorea-acanthocytosis, Cohen syndrome, early-onset Parkinson’s disease and spastic ataxia. Recent evidence suggests that the different VPS13 paralogs transport lipids between organelles at different membrane contact sites. How each VPS13 isoform is targeted to organelles is not known. We have shown that the localization of yeast Vps13 protein to membranes requires a conserved six-repeat region, the Vps13 Adaptor Binding (VAB) domain, which binds to organelle-specific adaptors. Here, we use a systematic mutagenesis strategy to determine the role of each repeat in recognizing each known adaptor. Our results show that mutation of invariant asparagines in repeats 1 and 6 strongly impact the binding all adaptors and block Vps13 membrane recruitment. However, we find that repeats 5 to 6 are sufficient for localization and interaction with adaptors. This supports a model where a single adaptor binding site is found in the last two repeats of the VAB domain, while VAB domain repeat 1 may help maintain domain conformation. Importantly, a disease-causing mutation in VPS13D, which maps to the highly conserved asparagine residue in repeat 6, blocks adaptor binding and Vps13 membrane recruitment when modeled in yeast. Our findings are consistent with a conserved adaptor binding role for the VAB domain and suggests the presence of as-yet-unidentified adaptors in both yeast and humans.

## Introduction

Membrane contact sites (MCSs) — regions where organelle membranes are tethered together in close proximity— are key sites of non-vesicular lipid and ion transport (1, 2). Dysfunction at these sites can perturb cellular functions including Ca^2+^ signalling, lipid transport, autophagy, intracellular protein transport and ER homeostasis (3–5). Indeed, a growing number of mutations in MCS proteins have been found to cause disease (3, 5).

A key example of disease-associated MCS proteins are the VPS13 paralogs, which were recently implicated in lipid transport at MCSs (6). Autosomal recessive mutations in the four human VPS13 proteins (VPS13A-D) are associated with distinct neurological disorders. Mutations in *VPS13A* cause chorea-acanthocytosis (OMIM 200150), a progressive movement disorder with abnormal red blood cell morphology (7). Mutations in *VPS13B* result in Cohen syndrome (OMIM 216550), a congenital multisystem disorder characterized by intellectual disability and developmental delay, and are implicated in autism (8–10). More recently mutations have been identified in *VPS13C* and *VPS13D* that respectively result in rapidly progressive, early-onset Parkinson’s disease (OMIM 616840) (11), and spastic ataxia (OMIM 607317) (12, 13). Disease-causing mutations in *VPS13A-C* are frequently null or truncating, resulting in no detectable protein (11, 14, 15), whereas *VPS13D* is essential and thus at least one allele retains some VPS13D functions in affected individuals (12, 13).

As VPS13 proteins are ubiquitously expressed, it is unclear why mutations in each isoform result in distinct diseases. Human VPS13 proteins may have diverged to function at different subsets of MCSs. Indeed, VPS13A and VPS13C localize to contact sites that link the endoplasmic reticulum (ER) to mitochondria and endosomes respectively (6), while VPS13B localizes to the Golgi and to endosomal vesicles (16, 17). VPS13A and VPS13C are also found at ER-lipid droplet contact sites suggesting they may have both distinct and overlapping roles (6, 18). Delineating the sites at which the different human paralogs function will provide a better understanding of how defects in lipid transport at specific MCSs contribute to neurological disorders.

Because VPS13 is conserved in eukaryotes (19), much of our understanding of these proteins arises from studies performed in the budding yeast, *Saccharomyces cerevisiae*. Yeast has a single Vps13 protein that localizes to multiple organelle membranes and MCSs (20–24). Structural studies show that the highly conserved Vps13 N-terminus forms a hydrophobic groove and is sufficient for lipid transport between membranes *in vitro* (6). This lipid transport function could explain why Vps13 can compensate for defects in the ER-mitochondrial encounter structure (ERMES), a lipid-transferring tether that forms ER-mitochondrial contact sites (21, 23). Other phenotypes of *vps13* mutants, such as aberrant actin morphology and Golgi-endosome trafficking defects (25, 26), are not clearly linked to lipid transport although this may be because the responsible MCSs have not yet been identified.

Understanding how VPS13 proteins are targeted to membranes could be key to understanding their role at different MCSs. We and others have recently demonstrated that specific adaptor proteins competitively target yeast Vps13 to different organelles (22, 24, 27) at different developmental stages or in response to environmental cues. The PX-domain protein Ypt35 recruits Vps13 to endosomes during exponential growth, while under starvation conditions, Ypt35 recruits Vps13 to the vacuole membrane at sites of contact with the nuclear ER (24). The mitochondrial adaptor, Mcp1, localizes Vps13 to the outer mitochondrial membrane (22, 24) whereas during meiosis, the sporulation-specific protein Spo71 recruits Vps13 to the prospore membrane (27).

These adaptors all possess a consensus P×P motif that interacts with Vps13 through a conserved domain found only in VPS13 proteins, which we have termed the Vps13 Adaptor Binding (VAB) domain (24). While the function of the VAB domain in human VPS13 proteins has not been determined, a VAB domain-containing fragment of VPS13C is sufficient for targeting to endolysosomal membranes suggesting that this domain may be important for the localization of at least some human VPS13 proteins (6). Importantly, disease-causing missense mutations in VPS13B and VPS13D alter conserved asparagine residues in the VAB domain (13, 28), suggesting it may have a conserved, functionally important role in yeast and humans.

The VAB domain consists of six repeats of approximately 100 residues with a highly conserved asparagine within the N-terminus of each repeat. It is not known if different adaptors bind different repeats and compete through steric hinderance, or if instead the VAB domain contains a single binding pocket that binds all motifs. In this study, we systematically mutate each of the yeast VAB domain repeats and demonstrate that the binding pocket for all known adaptors lies within repeats 5 to 6. We further demonstrate that a pathogenic missense mutation within the 6^th^ VAB domain repeat of VPS13D blocks Vps13 adaptor binding and recruitment to membranes when modeled in yeast, suggesting the binding of VPS13D to adaptor proteins may be relevant for disease.

## Results

### Specific VAB domain mutations disrupt binding of the endosomal adaptor Ypt35

To determine the contribution of each repeat to adaptor binding, we generated a series of Vps13 point mutants with alanine substitutions in the highly conserved asparagine of each repeat (24) (Fig 1A). Herein these mutants are referred to as Vps13^N1^, Vps13^N2^, and so on, indicating the repeat with the substitution.

**Figure 1:**
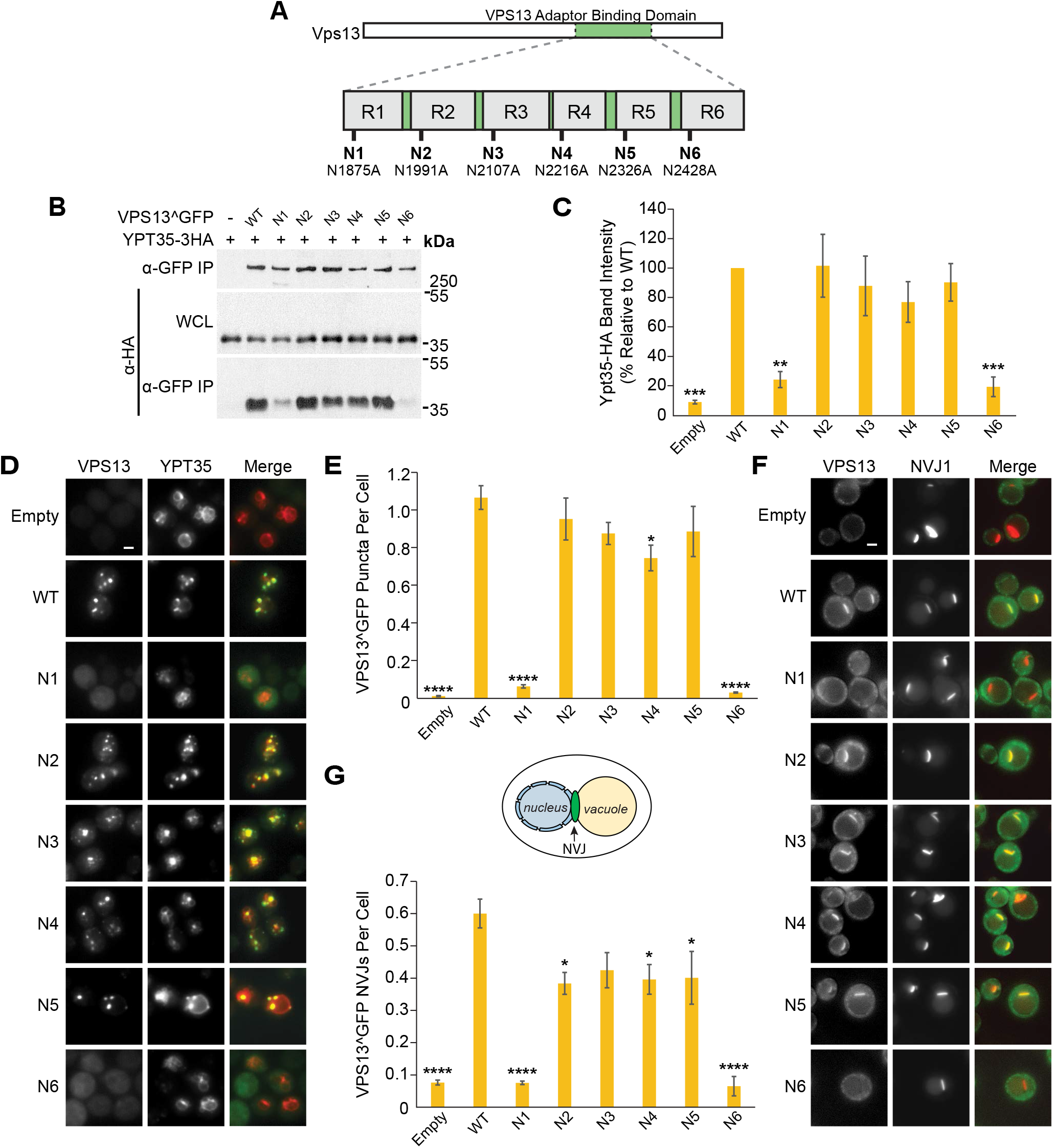
Mutations in the Vps13 VAB domain disrupt its interaction with the endosomal adaptor Ypt35. (A) Schematic of Vps13 VAB domain indicating the position of the mutated conserved asparagines in repeats 1 to 6 (R1-R6). (B) Coimmunoprecipitation of WT or mutant Vps13^GFP with Ypt35-3HA overexpressed from the *ADH1* promoter. Proteins were expressed on plasmids in *vps13Δ* strains. IP, immunoprecipitation; WCL, whole-cell lysate. (C) Densitometry analysis of Ypt35-HA co-immunoprecipitation bands shown in (B); one-way ANOVA with Dunneti-corrected post-hoc test; n=4; ***, P <0.001; **, P<0.01. (D) Localization of WT or mutant Vps13^GFP to bright puncta colocalizing with *ADH1pr-*driven Ypt35-RFP. Proteins were expressed on plasmids in *ypt35Δ* strains. (E) Automated quantitation of Vps13^GFP puncta from (D). One-way ANOVA with Dunneti-corrected post-hoc test; n=3; cells/strain/replicate ≥ 1370; ****, P <0.0001; *, P <0.05. (F) Localization of WT or mutant Vps13^GFP to NVJ1-RFP labelled NVJs in acetate-based media. Proteins were expressed on plasmids in *vps13Δ* strains. (G) Automated quantitation of the number of Vps13^GFP-labeled NVJs, from (F). The diagram shows the location of the NVJ. One-way ANOVA with Dunnett-corrected post-hoc test; n=3, cells/strain/replicate ⁥ 495; ****, P <0.0001; *, P <0.05. Error bars indicate SEM. Bars, 2 μm.

Each mutant was tagged with GFP at an internal site that does not disrupt known Vps13 functions (Vps13^GFP) (21) and expressed from a plasmid under its native promoter in a *vps13Δ* strain that co-expressed high levels of the HA-tagged endosomal adaptor, Ypt35. Mutation of the invariant asparagines did not alter protein levels substantially (Fig S1A), and mutations in repeats 2-5 did not significantly affect the co-immunoprecipitation (co-IP) of Ypt35 with Vps13 (Fig 1B,C). However, the Vps13^N1^ and Vps13^N6^ mutants demonstrated strongly reduced pull-down of Ypt35, with respective recoveries of 24.2% (P<0.01) and 19.3% (P<0.001) relative to WT.

We used fluorescence microscopy to determine the impact of each mutation on the Ypt35-mediated localization of Vps13^GFP to endosomes during exponential growth. In *ypt35Δ* strains expressing high levels of RFP-tagged Ypt35, wild type Vps13^GFP and most mutant forms localized to bright endosomal puncta that colocalized with Ypt35 (Fig 1D). Consistent with the decreased binding observed by co-IP, automated quantitation showed a near complete loss of localization of Vps13^N1^ and Vps13^N6^ mutants to puncta (Fig 1E; P<0.0001). A slight reduction in the punctate localization of Vps13^N4^ was also seen (69.8% relative to WT; P<0.05), although no significant loss of binding was detected by co-IP (Fig 1C).

To examine if the Ypt35-mediated recruitment of Vps13 to nucleus-vacuole junctions (NVJs) was similarly affected, we analysed the localization of Vps13^GFP mutants in *vps13Δ* strains expressing the NVJ marker Nvj1-RFP. Upon glucose starvation, the Vps13^N1^ and Vps13^N6^ mutants remained cytosolic and did not colocalize with Nvj1-RFP (Fig 1F,G; P<0.0001). A minor defect was observed for Vps13^N2^, Vps13^N4^ and Vps13^N5^, which localized to ~33% fewer NVJs compared to wildtype (P<0.05) (Fig 1G). Taken together, these results suggest that the conserved asparagines of the 1^st^ and 6^th^ VAB domain repeats are necessary for the Vps13-Ypt35 interaction.

### Similar VAB domain mutations disrupt the Vps13-Mcp1 interaction

To determine if different VAB repeats are required for the interaction of Vps13 with the mitochondrial adaptor Mcp1, the Vps13^GFP mutants were immunoprecipitated from *vps13Δ* strains expressing high levels of HA-tagged Mcp1 (Fig 2A). As with Ypt35, mutation of the first and sixth repeats dramatically reduced the interaction with Mcp1 (24.2% and 6.0% recovery relative to WT respectively; P<0.0001). Modest reductions in Mcp1 recovery were seen for Vps13^N3^ (66.8%, P<0.05) and Vps13^N4^ (42.4%, P<0.0001) (Fig 2B).

**Figure 2:**
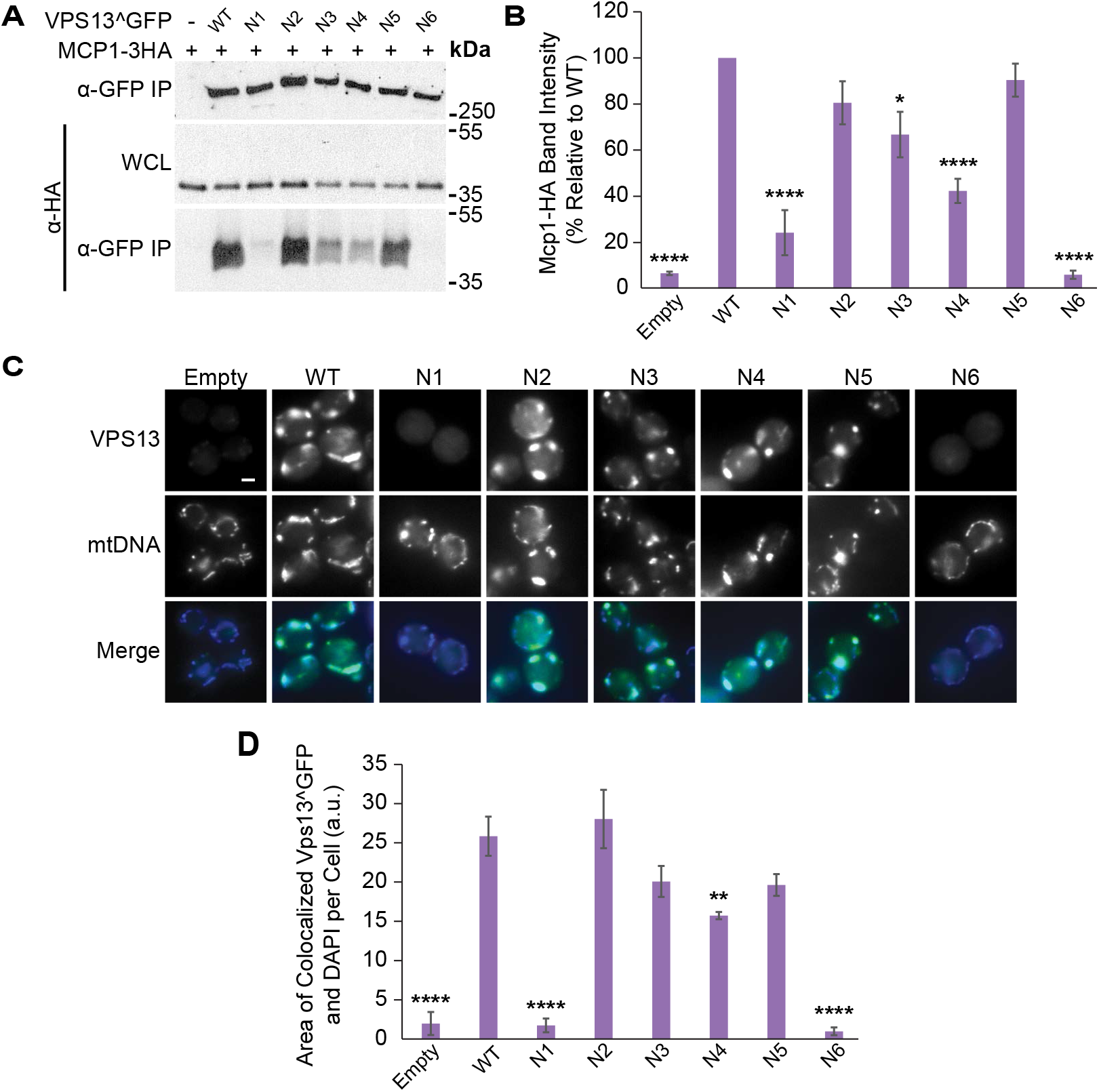
VAB domain mutations disrupt Vps13 Mcp1-mediated mitochondrial localization. (A) Coimmunoprecipitation of *ADH1pr*-driven Mcp1-3HA with WT or mutant Vps13^GFP. IP, immunoprecipitation; WCL, whole-cell lysate. (B) Densitometry analysis of Mcp1-HA co-immunoprecipitation bands shown in (A); one-way ANOVA with Dunneti-corrected post-hoc test; n=4; ****, P <0.0001; *, P<0.05. (C) Colocalization of WT or mutant Vps13^GFP and DAPI stained mtDNA in cells overexpressing Mcp1-3HA from an *ADH1* promoter. (D) Automated quantitation of colocalized Vps13^GFP and mtDNA area/cell from (C). One-way ANOVA with Dunneti-corrected post-hoc test; n=3, cells/strain/replicate ≥ 1012; ****, P <0.0001; **, P<0.01. Error bars indicate SEM. Bar, 2 μm.

The recruitment of the Vps13^GFP mutants to mitochondria was examined in the strains overexpressing Mcp1, which enhances Vps13 mitochondrial localization (22, 24) (Fig 2C). Automated analysis of the colocalization between Vps13^GFP and DAPI-stained mitochondrial DNA showed that Vps13^N1^ and Vps13^N6^ were defective in localizing to mitochondria (6.7% and 3.8% relative to WT; P<0.0001) (Fig 2D). In contrast, the mitochondrial localization of Vps13^N4^ was only slightly reduced (60.8% of WT) and that of Vps13^N3^ was not significantly altered, suggesting the modest reduction in Mcp1 binding was not sufficient to block Vps13 recruitment. Collectively, these results show that mutations in VAB domain repeats 1 and 6 strongly reduce binding of two different adaptors, Mcp1 and Ypt35.

### All three adaptors have similar requirements for VAB domain binding

A similar strategy was used to determine if the interaction of Vps13 with the prospore membrane adaptor, Spo71, required the same VAB domain repeats as Ypt35 and Mcp1. When Vps13^GFP mutants were immunoprecipitated in *vps13Δ* strains expressing high levels of the HA-tagged Spo71 (Fig 3A), only Vps13^N1^ and Vps13^N6^ had reduced pull-down of Spo71 (40.7% and 14.0% relative to WT; P<0.05 and P<0.01 respectively) (Fig 3B).

**Figure 3:**
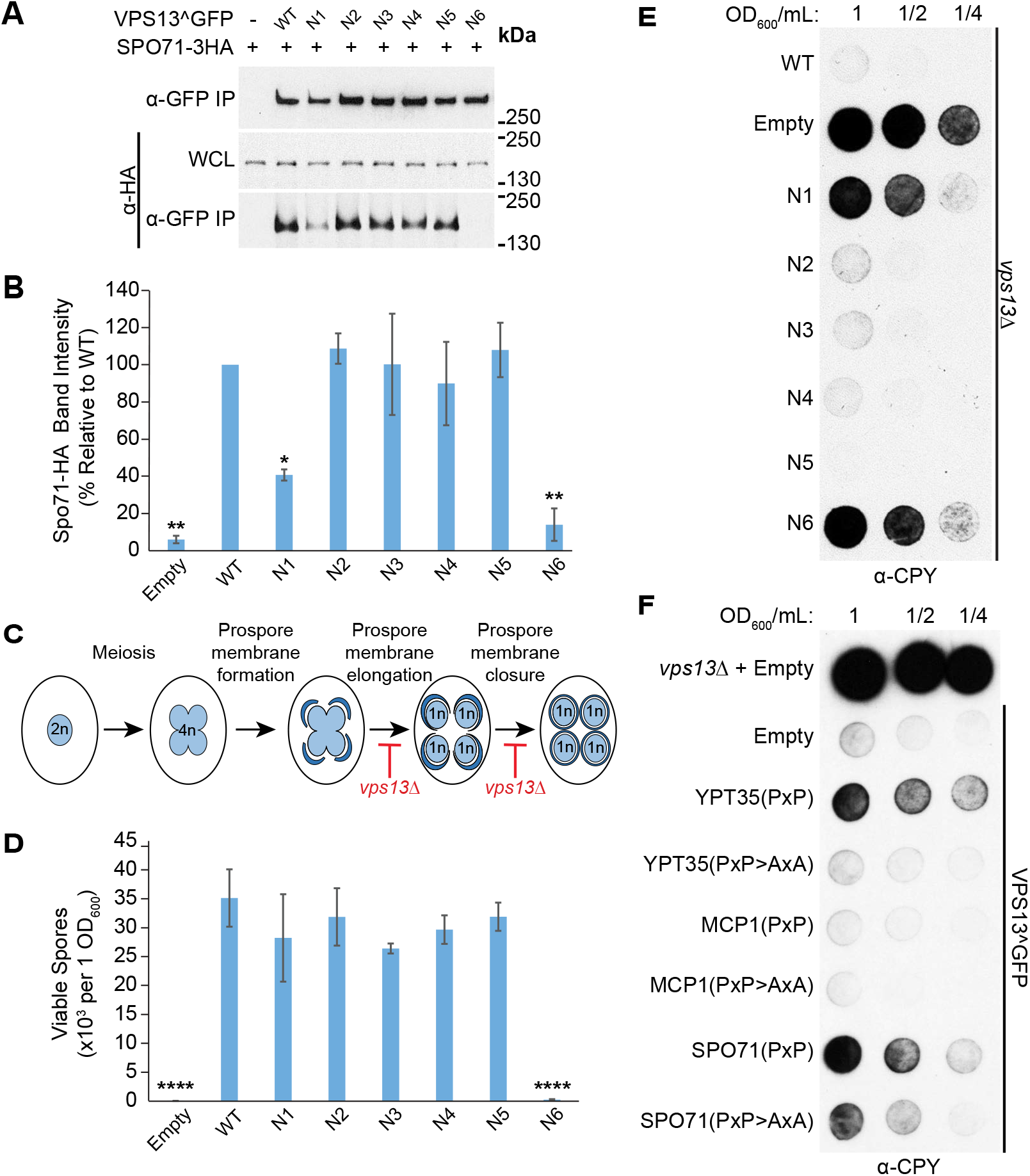
VAB domain mutations disrupt Spo71-mediated sporulation and CPY sorting. (A) Coimmunoprecipitation of WT or mutant Vps13^GFP with Spo71-3HA overexpressed from a *TEF1* promoter. IP, immunoprecipitation; WCL, whole-cell lysate. (B) Densitometry analysis of Spo71-HA co-immunoprecipitation bands shown in (A); one-way ANOVA with Dunneti-corrected post-hoc test; n=3; **, P <0.01; *, P <0.05. (C) Model of yeast sporulation depicting prospore membrane formation, elongation and closure. Red bars indicate steps in sporulation that are defective in *vps13Δ* strains. (D) Mutation in the sixth VAB domain repeat prevents viable spore formation. Sporulated *vps13Δ* diploids carrying WT or mutant VPS13^GFP on plasmids were plated on selective media and colonies counted atier 3 days of growth at 30 °C. One-way ANOVA with Dunneti-corrected post-hoc test; n=3; ****, P<0.0001. (E) Mutations in the VAB domain result in CPY secretion. *vps13Δ* strains carrying WT or mutant VPS13^GFP on plasmids were spotied at the indicated dilutions, covered in nitrocellulose, grown overnight at 30 °C and blotied with α-CPY antibody to detect secretion. Representative image shown; n=3. (F) Overexpression of adaptor P×P-motifs results in CPY secretion that is rescued by mutating the conserved prolines within the motifs. The following fragments were expressed as RFP fusions from the strong *TEF1* promoter: Ypt35(1-48), Mcp1(1-58) and Spo71(359-411) with P×P>AxA fragments containing Ypt35(P10,12A), Mcp1(P9,11A) and Spo71(P390,392A) substitutions. Error bars indicate SEM. Representative image shown; n = 3.

During sporulation, Vps13 is recruited by Spo71 to the prospore membrane which elongates and closes around haploid daughter nuclei (27). *vps13Δ* diploids have defects in prospore membrane expansion and closure preventing the formation of viable haploid gametes (Fig 3C)(20). To examine the phenotypic consequences of the reduced Vps13-Spo71 interaction, we determined the sporulation efficiency of *vps13Δ* diploids expressing each VAB domain mutant. All mutants gave rise to a similar number of viable haploids relative to wildtype, except for Vps13^N6^ which was nearly unable to form viable spores (P<0.0001) (Fig 3D).

We were surprised that the Vps13^N1^ mutant, which showed a 60% reduction in Spo71 binding in our pulldown assay, did not have a sporulation defect. This suggests the level of Spo71 binding provided by Vps13^N1^ meets the threshold needed for wild type function. Taken together, our systematic mutational analysis of the Vps13 VAB domain shows that the binding of all adaptors is strongly affected by mutations in repeat 1 and 6, with repeat 6 being the most critical for adaptor interaction.

### VAB domain mutations result in CPY secretion

Vps13 was initially identified in a screen for mutants that missort the vacuolar hydrolase carboxypeptidase Y (CPY) (29). In *vps13*Δ cells, defects in recycling the CPY receptor between the Golgi and endosomes causes CPY to enter the secretory pathway and be released from the cell. Because the known adaptors are not required for this process, we tested the effect of the VAB domain mutations on CPY sorting. Vps13^GFP mutants were spotted on plates and covered with a nitrocellulose filter to capture secreted CPY, which was detected by western blotting (Fig 3E). CPY was secreted by both Vps13^N1^ and Vps13^N6^ mutants, suggesting that the first and sixth repeats are necessary for the CPY sorting function of Vps13. This could indicate that a novel Vps13 adaptor is required for CPY sorting.

We have previously shown that soluble chimeric proteins carrying a P×P motif compete with adaptors to block Vps13 recruitment to membranes (24). Therefore, if a new adaptor is required for Vps13’s CPY sorting function, overexpression of soluble P×P motif proteins should cause cells expressing wild type Vps13 to secrete CPY. Indeed, when the Ypt35 and Spo71 P×P motifs, but not the weaker Mcp1 motif, were fused to a cytosolic RFP marker and expressed from the strong *TEF1* promoter in Vps13^GFP strains, a clear CPY sorting defect was observed that required the conserved prolines (Fig 3F). This suggests that Vps13 may be recruited by an as-yet-undiscovered P×P motif-containing adaptor to membranes where it functions in the CPY sorting pathway.

### VAB repeats 5-6 are sufficient for adaptor binding

Our results suggest that the integrity of the first and sixth repeats of the VAB domain are critical for binding all adaptor proteins. To determine if these repeats are likely to form a single binding pocket in the folded protein, we used I-TASSER (30) to generate a predicted VAB domain structure from yeast and human VPS13 proteins. The highest-scoring models suggest the VAB domains of VPS13A, VPS13B, VPS13C and yeast Vps13 form two six-bladed β-propellers (Fig S2). In these models, each repeat forms 2 blades of a propeller with the conserved asparagines residing in alternate blades. This region of the human VPS13A and VPS13C was previously suggested to have a WD40-like fold (6); however, because Vps13 lacks the conserved “WD” residues that typically characterize WD40 proteins (31, 32), we will use the more general term of β-propeller.

To identify the minimal subdomain required for adaptor binding, we expressed Envy-tagged forms of each predicted VAB β-propeller from the *ADH1* promoter in strains overexpressing HA-tagged Ypt35 (Fig 4A). Both predicted β-propeller domains were stably expressed but only the second β-propeller, corresponding to repeats 4-6, bound Ypt35. This fragment pulled down all three adaptors in a P×P motif-dependent manner (Fig S1B-D). Some β-propeller domains can be truncated into stable fragments corresponding to one or more blades (33, 34). Further truncations of the second predicted β-propeller determined that repeats 5-6, but not repeats 5 or 6 alone, could pull down Ypt35 (Fig 4B). Indeed, repeats 5-6 were sufficient for the P×P motif-dependent interaction of all 3 adaptors (Fig 4C-E).

**Figure 4:**
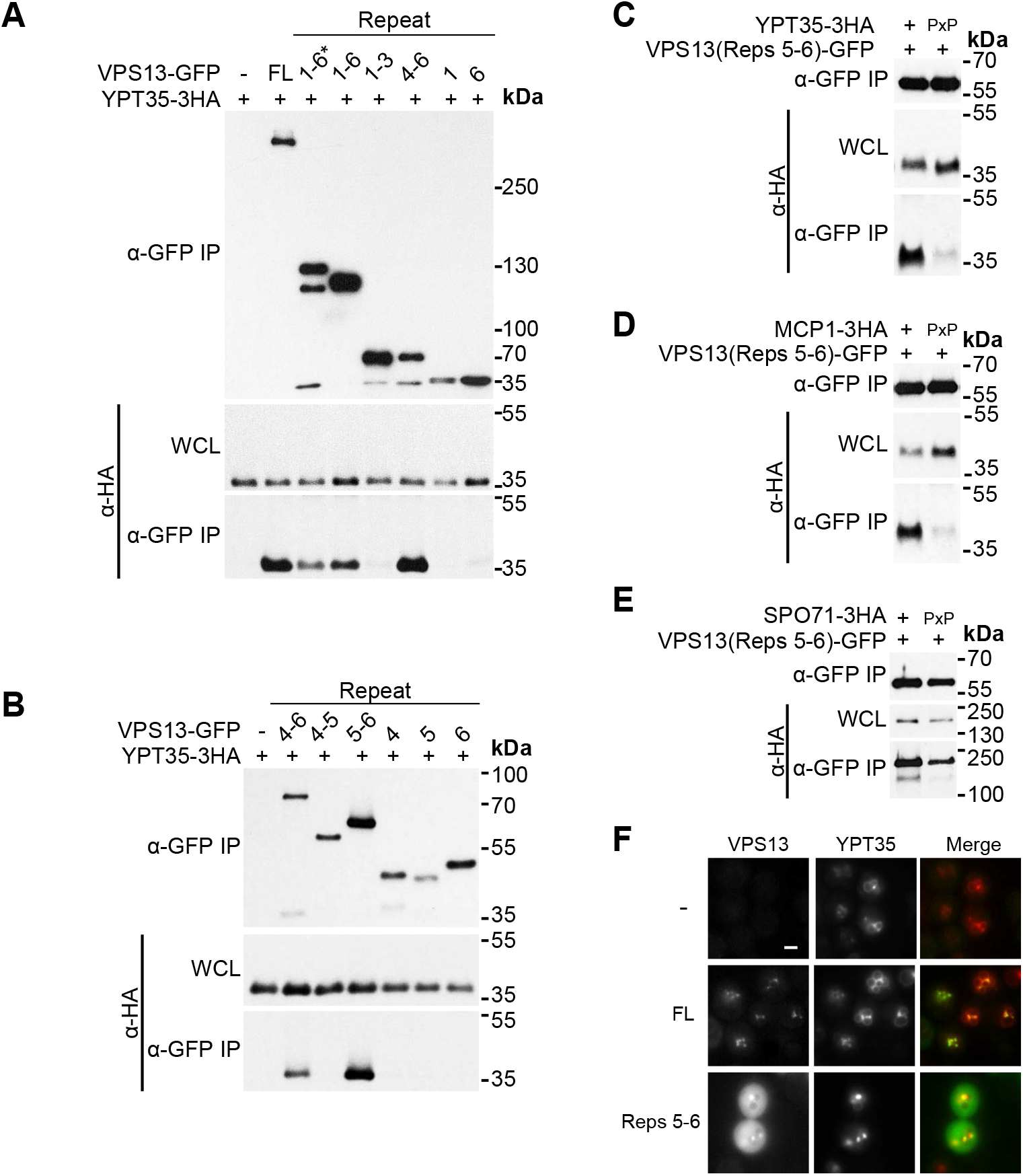
VAB domain repeats 5-6 are sufficient for binding all three adaptors. (A) VAB domain repeats 4-6 are sufficient for interaction with Ypt35. Coimmunoprecipitation of *ADH1pr*-driven Ypt35-3HA with ENVY-tagged Vps13 VAB domain truncations. 1-6* indicates an N-terminally extended VAB domain sequence. (B) VAB domain repeats 5-6 are sufficient for interaction with Ypt35. Coimmunoprecipitations of Ypt35-3HA with ENVY-tagged Vps13 VAB domain repeats 4-6 truncations. (C-E) Adaptor interaction with VAB domain repeats 5-6 is P×P motif dependent. Coimmunoprecipitations of WT or P×P motif mutant overexpressed Ypt35-3HA, Mcp1-3HA and Spo71-3HA with ENVY-tagged Vps13 VAB repeats 5-6. IP, immunoprecipitation; WCL, whole-cell lysate. (F) Vps13 VAB repeats 5-6 are sufficient for colocalization with *ADH1pr*-driven Ypt35-RFP. Representative images selected from n = 3. Bar, 2 μm.

To determine if repeats 5-6 were sufficient for recruitment to membranes, we examined the localization of this fragment in *ypt35*Δ cells expressing high levels of RFP-tagged Ypt35 (Fig 4F). Indeed, GFP-tagged VAB repeats 5-6 colocalized with Ypt35-RFP labelled puncta, although fewer puncta were formed compared to full-length Vps13^GFP.

Collectively, these results indicate that the adaptor binding pocket of the Vps13 VAB domain is located in repeats 5-6, within the second predicted β-propeller domain. This is consistent with our mutational analysis which showed mutation of the invariant asparagine in repeat 6 severely reduced the binding of all three adaptors. In contrast, the conserved asparagine of repeat 1, which is predicted to lie within the first β-propeller, appears to be important for binding only in the context of the full-length protein.

### Human disease-causing mutations disrupt adaptor binding in yeast Vps13

Though the VAB domains of human VPS13 proteins lack known binding partners, disease-causing mutations in the VAB domains of VPS13B and VPS13D have been identified (Fig 5A). In VPS13D, a serine substitution in the conserved asparagine of VAB domain repeat 6, N3521S, is causative for spastic ataxia (13). In VPS13B, a serine substitution of the conserved asparagine of VAB domain repeat 4, N2993S, was identified as a homozygous mutation in a Belgian sibling pair presenting with Cohen syndrome (28).

**Figure 5:**
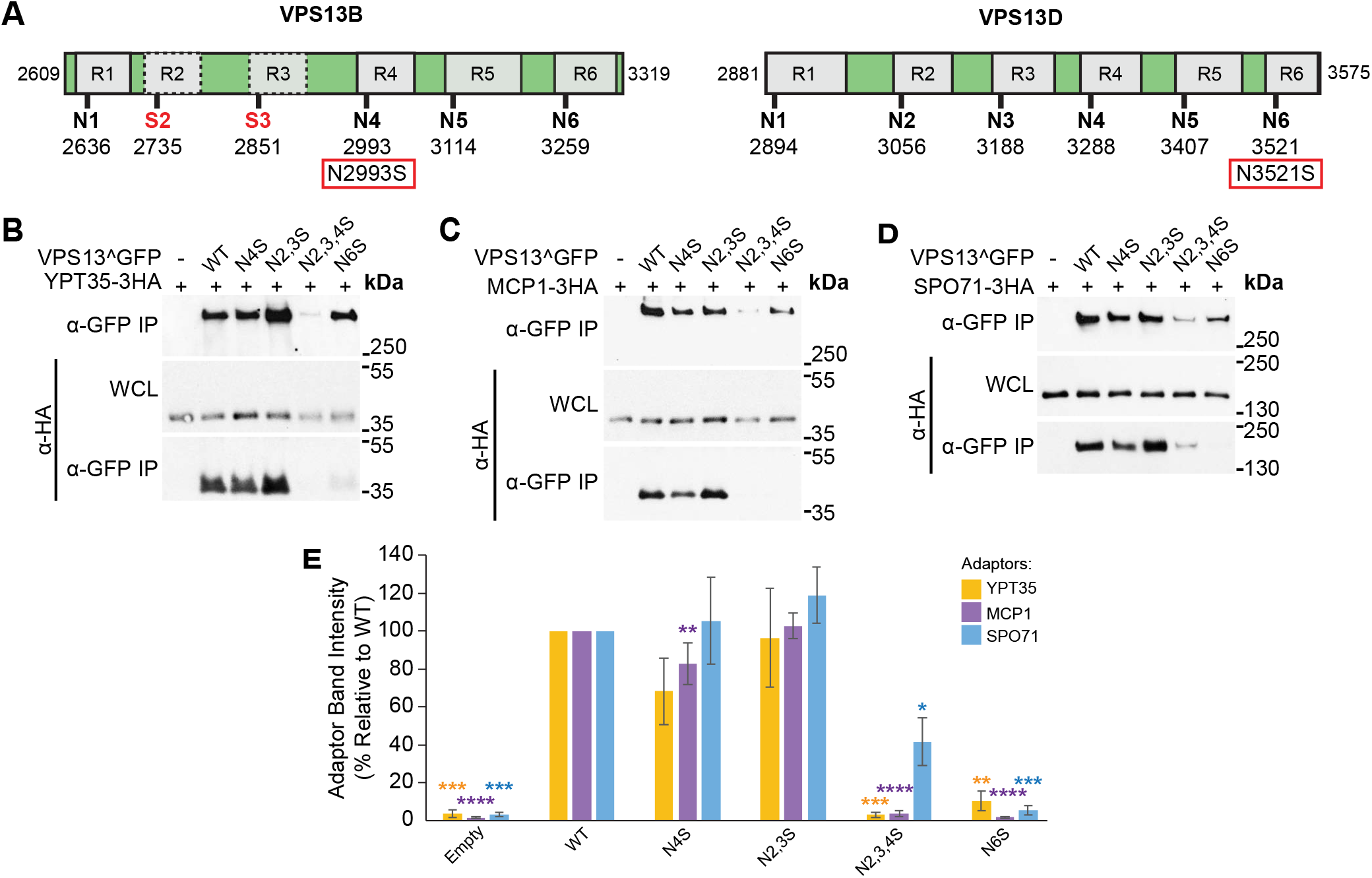
Causative Cohen syndrome and spastic ataxia mutations result in perturbed adaptor binding in yeast. (A) Schematic of the VAB domains of VPS13B and VPS13D indicating the six repeats and position of conserved asparagines. VPS13B has a divergent VAB domain with serines in place of asparagines in repeats 2 and 3 indicated by red text. The causative Cohen syndrome mutation in VPS13B and spastic ataxia mutation in VPS13D are indicated in red boxes. (B-D) Coimmunoprecipitations of overexpressed Ypt35-3HA, Mcp1-3HA and Spo71-3HA with WT or mutant Vps13^GFP modeling the human mutations. IP, immunoprecipitation; WCL, whole-cell lysate. (E) Densitometry of immunoprecipitated HA-tagged adaptors from (B-D). One-way ANOVA with Dunneti-corrected post-hoc test; n = 3; ****, P<0.0001; ***, P<0.001; **, P<0.01; *, P<0.05. Error bars indicate SEM.

We modeled the VPS13D spastic ataxia mutation by mutating the corresponding N2428 residue in yeast Vps13^GFP to produce Vps13^N6S^ and tested the effect on adaptor binding in a pull-down assay. The Vps13^N6S^ mutant showed background level of binding to all three adaptors (n=3 per adaptor; Ypt35 = P<0.01; Mcp1 = P<0.0001; Spo71 = P<0.001), similar to the Vps13^N6^ mutant where the same residue was substituted with alanine (Fig 5B-E). This suggests the disease-causing mutation in VPS13D disrupts the function of the VAB domain in the context of yeast Vps13.

We next modeled the pathogenic VPS13B N2993S mutation in yeast by mutating the corresponding N2216 residue in Vps13^GFP to create Vps14^N4S^. Vps13^N4S^ had a small but significant reduction in the pull-down of Mcp1 (76.9% relative to wildtype; P<0.01), but not the other two adaptors (Fig 5B-E), consistent with the phenotype of the Vps13^N4^ substitution (see Fig 2B). VPS13B has the least conserved VAB domain of the human paralogs, with serines replacing the conserved asparagines in the 2^nd^ and 3^rd^ repeats. When these changes were modeled in yeast, creating Vps13^N2,3S^, we saw no effect on protein stability or adaptor pull-down (Fig 5B-E) suggesting these substitutions do not grossly disrupt the structural integrity of the VAB domain. However, more dramatic defects were seen when a N4S mutation was made in the context of this divergent VAB domain. The Vps13^N2,3,4S^ mutant showed a strongly decreased protein level, which likely accounts for the reduced pulldown of each adaptor (Fig 5B-D). While Ypt35 and Mcp1 were recovered at background levels (P<0.001 and P<0.0001 respectively), recovery of Spo71 was 41.6% relative to wildtype (P<0.05), suggesting the residual Vps13^N2,3,4S^ was still capable of binding Spo71.

These Vps13^GFP mutants were also examined for their ability to localize to organelles and mediate CPY sorting. Vps13^N2,3,4S^ and Vps13^N6S^ were not recruited to bright puncta by Ypt35-RFP (Fig 6A-B)(P<0.0001), or to mitochondria in cells expressing high levels of Mcp1 (Fig 6C-D) (P<0.01). Both Vps13^N2,3,4S^ and Vps13^N6S^ missorted CPY (Fig 6E), whereas no defect was seen for the Vps13^N2,3S^ or Vps13^N4S^. Together these results suggest that VPS13B and VPS13D disease-causing mutations within the VAB domain result in reduced protein function when modeled in yeast, but for different reasons. Whereas the VPS13B mutation disrupts protein stability, the pathogenic VPS13D mutation does not significantly change protein levels, but instead may block its ability to interact with binding partners.

**Figure 6:**
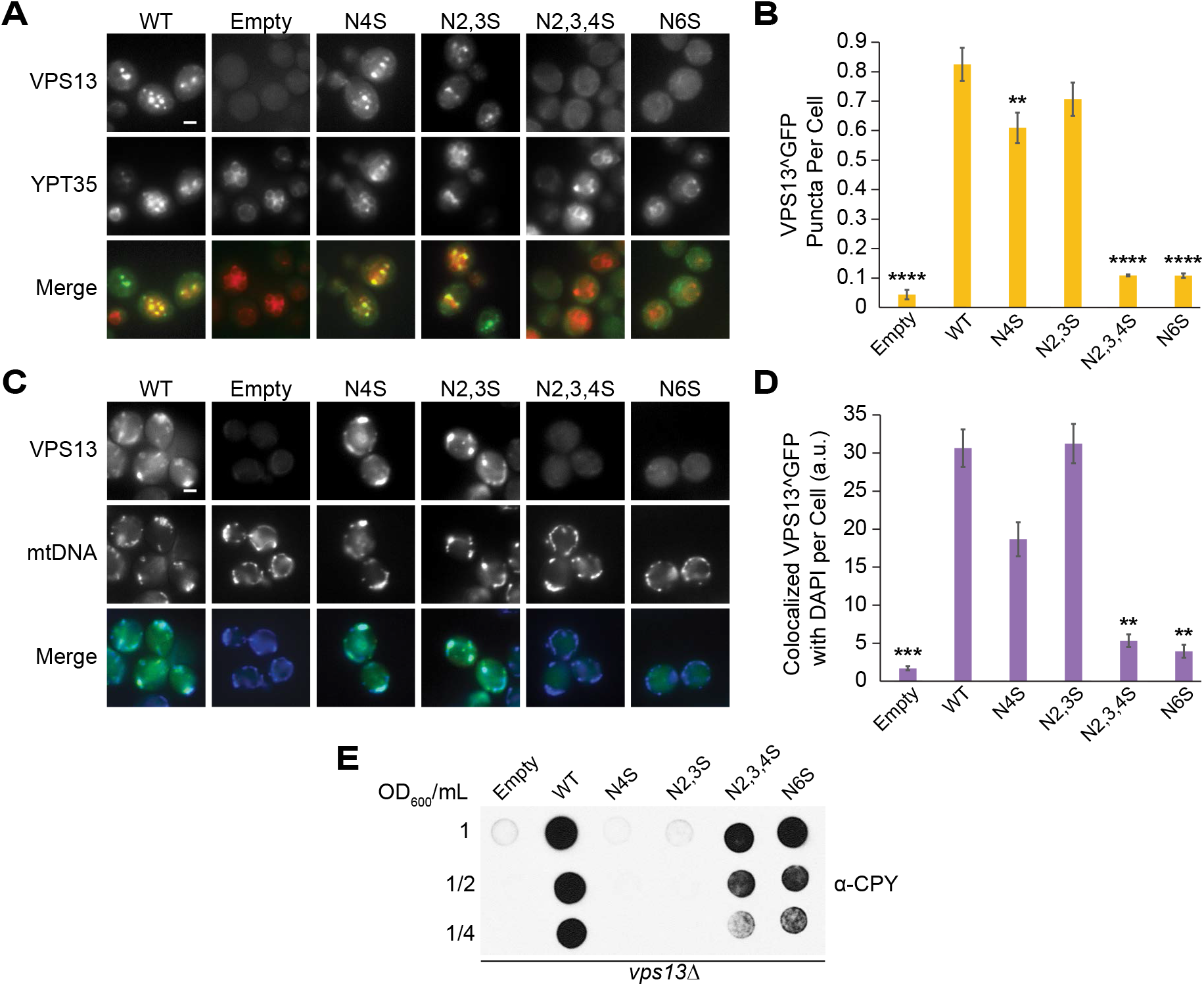
Causative Cohen syndrome and spastic ataxia mutations result in loss of yeast Vps13 localization and CPY secretion. (A) Disease-causing mutations in the Vps13 VAB domain block recruitment to *ADH1pr*-driven Ypt35-RFP. Proteins were expressed on plasmids in *ypt35Δ* cells. (B) Automated quantitation of Vps13^GFP puncta from (A). One-way ANOVA with Dunneti-corrected post-hoc test; n=3, cells/strain/replicate ≥ 1164; ****, P <0.0001; **, P <0.01. (C) Disease-causing mutations in the Vps13 VAB domain block recruitment to mitochondria. Colocalization of Vps13^GFP mutants and DAPI stained mtDNA in cells overexpressing Mcp1-3HA from an *ADH1* promoter. (D) Automated quantitation of colocalized Vps13^GFP and mtDNA area/cell from (C). One-way ANOVA with Dunneti-corrected post-hoc test; n=3, cells/strain/replicate ≥ 1200; ***, P <0.001; **, P <0.01; *. (E) Human disease-causing mutations in the VAB domain result in CPY secretion. *vps13Δ* cells carrying WT or mutant VPS13^GFP were blotied with α-CPY antibody to detect secretion as described in Fig 3E. Error bars indicate SEM. Bar, 2 μm.

## Discussion

The VAB domain, which is found only in VPS13 proteins, is conserved in yeast and human isoforms suggesting it has an important functional role. Our studies of yeast Vps13 indicate that mutation of an invariant asparagine residue in the sixth repeat of this domain disrupts a critical protein interaction site. Because mutation of the corresponding residue in human VPS13D causes disease, both human and yeast VPS13 proteins may use this site to bind key regulators. This demonstrates that yeast Vps13 provides a useful model for studying the impact of human disease mutations.

### The VAB domain is predicted to form a β-propeller structure

We found all adaptors respond similarly to alteration of the VAB domain repeats: mutation of the 6^th^ repeat abolished the binding of all three adaptors tested, while mutation of the 1^st^ repeat caused a partial defect. Moreover, we found that repeats 5 and 6 alone bound strongly to all three adaptors, suggesting adaptors recognize a single site within these repeats, consistent with our previous finding that the adaptors compete to recruit Vps13 to different membranes (24).

Structural modeling algorithms predict the VAB domains from yeast and human proteins form WD40-like β-propeller structures. The highest confidence models suggest the VAB domain folds into two 6-bladed β-propellers with each repeat contributing 2 blades. In this model, the adaptor binding site is contained within 4 blades of the second β-propeller (Fig 7A). Because mutation of the conserved asparagine in the 6^th^ repeat nearly abolished adaptor binding, whereas mutation of the analogous residue in the 5^th^ repeat had no effect, it is tempting to conclude that the “N6” residue lies in or near the binding pocket. As β-propellers can bind peptides at many sites along the top, side, or bottom surface (32, 35), it is also possible the conserved asparagine does not interact directly with the P×P peptide, but instead helps maintain the proper conformation of the binding site by stabilizing blade interfaces or maintaining the position of loops.

**Figure 7:**
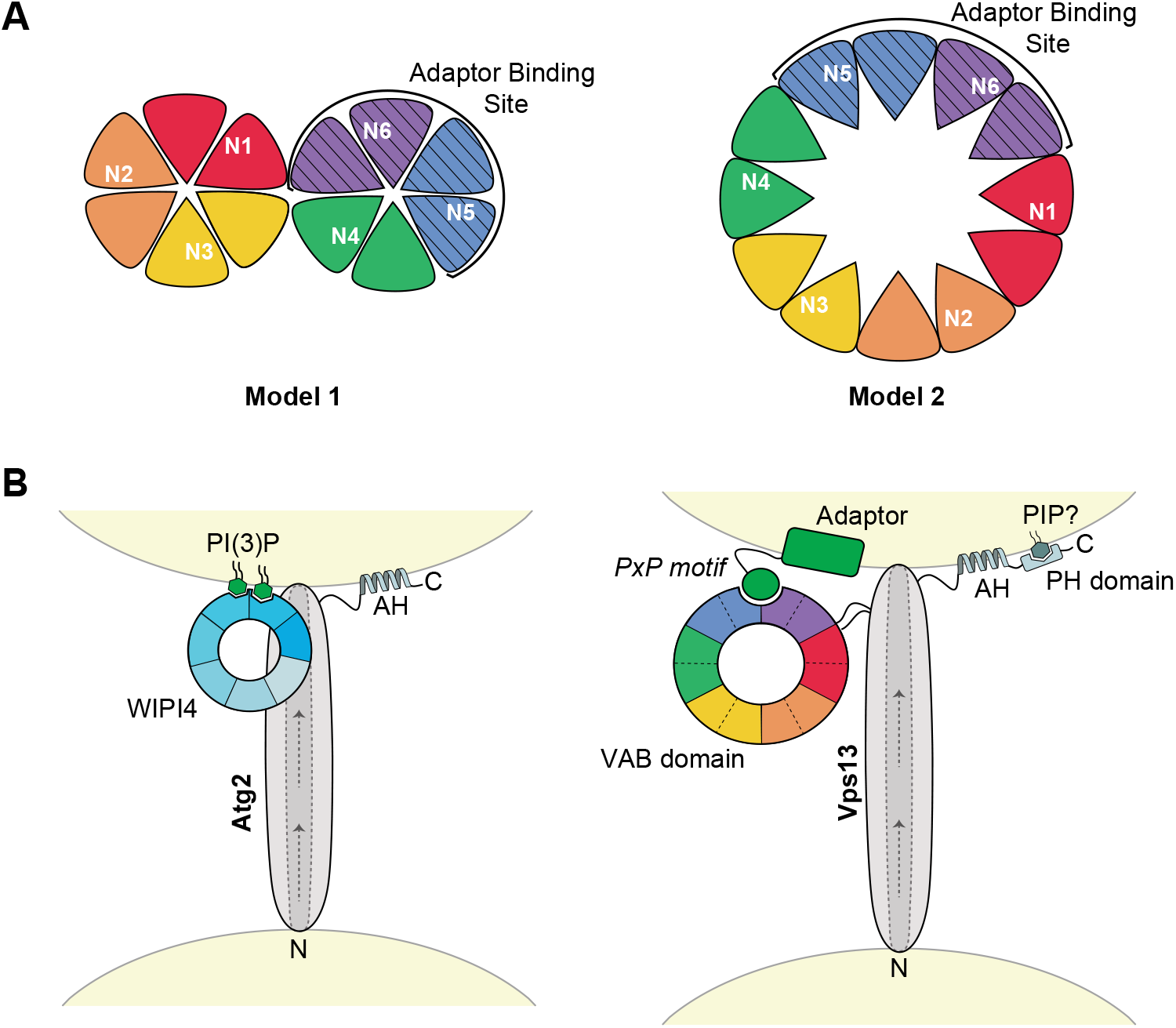
Model of the proposed VAB domain structure. (A) Proposed models of the VPS13 VAB domain with repeats 1 to 6 shown from red to purple. Each repeat contributes 2 blades to a β-propeller fold with the conserved asparagines in the first blade of each repeat. The adaptor-binding site, indicated by stripes, is within repeats 5-6. In model 1, the VAB domain forms two 6-bladed β-propellers and in model 2 this domain forms a single 12-bladed β-propeller. (B) Model of Atg2 and Vps13 recruitment to membranes through interaction with WIPI4 or the VAB domain respectively. In Atg2, the 7-bladed WD-40 protein, WIPI4, binds 2 molecules of PI(3)P at the isolation membrane via blades 5 and 6 (46, 47) and interacts with Atg2 via blade 2 (45) to target it to membranes. In Vps13, the VAB β-propeller structure binds P×P motif-containing adaptors through an interaction with the binding pocket in repeats 5-6. Other conserved C-terminal domains, such as amphipathic helices (AHs) and a predicted PH domain, may help position these proteins at membranes. Arrows indicate the putative direction of lipid transport along a hydrophobic groove shown by dotied lines. The N and C-termini of both proteins are indicated. Models are not to scale.

Mutation of the conserved asparagine in the 1^st^ VAB domain repeat also had a pronounced effect on adaptor binding. In the tandem β-propeller model, this repeat is located in the first β-propeller, which was dispensable for binding when the second β-propeller was expressed alone. In the intact Vps13 protein, the VAB domain may have a “clamshell” arrangement similar to other tandem β-propeller proteins where the first domain stabilizes the conformation of the second domain (36–38), or contributes a loop to a shared binding interface (39, 40). Alternatively, repeat 1 could contribute a β-strand to the final blade of repeat 6 in the second β-propeller to form a “molecular velcro”(36, 41, 42).

This tandem 6-bladed β-propeller model may be favoured by prediction algorithms because such structures are frequently found in structural databases (31, 32, 43). An alternate model is that the VAB domain forms a single 12-bladed β-propeller, an uncommon but previously solved structure (44) (Fig 7A). This model is consistent with single-particle EM images that suggest yeast Vps13 forms a rod-like structure with a large circular loop near the expected site of the VAB domain (26). In the single 12-bladed β-propeller model, repeats 1 and 6 are adjacent which could explain why mutation of the first repeat destabilizes a binding pocket within repeats 5-6.

This model places the adaptor binding site close to the membrane and to the Vps13 lipid-transporting stalk, while the remaining repeats project away from the lipid-transporting core where they could bind additional regulatory factors (Fig 7B). This arrangement is similar to that of the Vps13-related protein Atg2 and its interacting WD40 protein, WIPI4 (Atg18), which binds phosphatidylinositol 3-phosphate (PI(3)P) to enhance Atg2 recruitment to isolation membranes (45–47). Additional conserved C-terminal regions such as amphipathic helices in Atg2 (48, 49) and Vps13 (6), or the predicted PH domain in Vps13 (50), may help stabilize recruitment through coincidence detection. Structural studies will be needed to verify the predicted tandem 6-bladed or single 12-bladed β-propeller structure and identify the key residues involved in the Vps13-adaptor interaction.

### New adaptors in yeast and humans

Our work suggests the full complement of adaptor proteins that bind the yeast Vps13 VAB domain has not yet been uncovered and that an undiscovered P×P-containing adaptor likely regulates Vps13’s CPY sorting function. Such an adaptor may recruit Vps13 to Golgi membranes to function in Golgi/endosome transport and homotypic fusion (26). Vps13 has also been localized to other organelles, including peroxisomes (22), which could require yet other adaptors. These unidentified adaptors may be identified by bioinformatics approaches that identify predicted P×P motifs (24).

Novel adaptor proteins may also bind the VAB domains of human VPS13 proteins. The dramatic effect of the disease-causing VPS13D mutation when modeled in yeast strongly suggests that VPS13D has a conserved adaptor binding site. However, the P×P motif is not conserved beyond the order of *Saccharomycetales* suggesting the yeast P×P motif co-evolved with the VAB domain. It is interesting to speculate that the human VPS13 proteins have diverged to recognize distinct motifs that target them to discrete MCSs.

### The role of VAB domain in human VPS13 proteins

*VPS13D* is an essential gene and thus disease-causing mutations are typically compound heterozygous with one null and one missense VPS13D allele (12, 13). Because the VPS13D^N6S^ mutation is paired with a frame-shift mutation that removes nearly half of the protein sequence (13), it is likely that the VAB domain mutation maintains essential VPS13D functions while blocking a specific adaptor interaction that results in ataxia. Two other disease-causing missense mutations in VPS13D fall within the VAB domain in repeat 1 (L2900S) and repeat 3 (R3253N) (13). The effect of these mutations has not been studied, but the clustering of causative missense mutations in the VPS13D VAB domain is consistent with an important role in disease for this domain. Although the localization of VPS13D is unknown, the abnormal mitochondrial morphology seen in affected patient fibroblasts and in *Drosophila* models suggests that VPS13D may function at mitochondria (12, 51).

If human VPS13 proteins have evolved different localization mechanisms to target distinct contact sites, the VAB domains of some paralogs may have lost their adaptor binding function. This could be true for VPS13B, which has a poorly conserved VAB domain and instead localizes to Golgi membranes through its C-terminus (52). The divergent VAB domain of VPS13B has serine residues in the place of conserved asparagines in the 2^nd^ and 3^rd^ VAB domain repeats. Modeling these substitutions in yeast suggests they have little or no effect on protein function, and naturally occurring asparagine-to-serine substitutions in VAB repeats 3 and 5 are also present in members of the division *Magnoliaphyta* and *Ascomycota*, respectively. The observation that this divergent VAB domain is destabilized by an asparagine-to-serine substitution introduced into repeat 4 could explain why the N2993S mutation in VPS13B causes Cohen syndrome, which is typically characterized by a loss of VPS13B protein (15, 53). These results demonstrate the utility of the yeast system for modeling human disease mutations. Further studies in yeast are expected to generate new insights into membrane targeting mechanisms of the VPS13 family and will provide a useful system for testing the effects of candidate pathogenic mutations.

Do other VPS13 VAB domains function in protein localization? A fragment of VPS13C containing its VAB domain is sufficient for localization to endolysosomal membranes (6). A similar VPS13A fragment was found to be cytosolic, however a missense mutation in this domain results in chorea-acanthocytosis (54). This suggests that VPS13A may rely on the VAB domain for localization, although the relevant adaptors may only be present in certain cell types, developmental stages or conditions.

Other features of the VPS13 proteins may work together for targeting to membrane contact sites. VPS13A and VPS13C both have additional localization determinants including a FFAT motif for ER targeting and an amphipathic helix for targeting to lipid droplets (6). Additionally, all human VPS13 proteins have been found to bind Rab proteins in high and low-throughput studies (17, 52, 55). Other regions of Vps13 may interact with lipids to stabilize membrane recruitment through coincidence detection. Yeast Vps13 binds lipids through regions at its N and C-terminus, and a causative chorea-acanthocytosis mutation in the APT1 domain disrupts binding to PI(3)P when modeled in yeast (25, 26). Identifying the different targeting mechanisms used by each human VPS13 protein will be important for determining the subset of MCSs where each paralog functions and for unravelling the role of specific MCSs in disease.

## Materials and Methods

### Yeast Strains and Plasmids

Strains and plasmids used in this study are described in Table S1 (A-B). All strains from this study were made by PCR-based homologous recombination as described (56, 57). Gene deletions were confirmed by colony PCR. Plasmids were made by homologous recombination in yeast by co-transforming linearized plasmids with generated PCR products. The plasmids were recovered in *Escherichia coli* and sequenced. All primers used in the generation of PCR products for strain and plasmid construction are listed in Table S1 (C).

### Coimmunoprecipitations

For Vps13^GFP point mutants, yeast were grown to log-phase in minimal synthetic dextrose media and 60 OD_600_ of cells stored at −80° C until further use. Cells were thawed and resuspended in 500 μL lysis buffer (50 mM HEPES, 0.1% Tween-20, 50 mM NaCl, 1 mM EDTA, 1 mM PMSF, and 1× yeast/fungal ProteaseArrest, pH 7.4) and lysed by vortexing with 0.5 mm diameter glass beads. 50 μL of lysate was set aside and resuspended in 4× Laemmli Sample Buffer with 8M urea (8% SDS, 40% glycerol, 240 mM Tris-Cl pH 6.8, 0.004 % bromophenol blue, 20% β-mercaptoethanol). The remainder of the lysates were incubated with polyclonal rabbit anti-GFP (EU2; Eusera) for 1 hour followed by Protein A-Sepharose beads (GE Healthcare) for another hour at 4° C. Washed beads were resuspended in 50 μL of Thorner buffer (8 M urea, 5% SDS, 40 mM Tris pH 6.8, 0.1 M EDTA, 1% β-mercaptoethanol, 0.4 mg/mL bromophenol blue) and proteins eluted by heating at 100° C for 5 minutes. Samples were run on SDS-PAGE gels, transferred to nitrocellulose membranes and blotted with either polyclonal rabbit anti-GFP (EU2; Eusera) and Tidyblot Western Blot Detection Reagent:HRP (STAR209P; Bio-Rad) or monoclonal mouse anti-HA (MMS-101R; Covance) and horseradish peroxidase-conjugated polyclonal goat anti-mouse (115-035-146; Jackson ImmunoResearch Laboratories).

For coimmunoprecipitations with Vps13 VAB domain truncations, 50 OD_600_ of yeast were lysed. The nitrocellulose membranes were blotted with either monoclonal mouse anti-GFP (11-814-460-001; Roche) or monoclonal mouse anti-HA (MMS-101R; Covance) followed by horseradish peroxidase-conjugated polyclonal goat anti-mouse (115-035-146; Jackson ImmunoResearch Laboratories). All blots were developed with West Pico (Pierce) or Amersham ECL Prime (GE Healthcare) chemiluminescent reagents and exposed to Amersham Hyperfilm (GE Healthcare). Films were scanned, and densitometry performed in Fiji (58).

### Fluorescence Microscopy and Automated Quantitation

Yeast were grown to log-phase in minimal synthetic dextrose media, transferred to concanavalin A-treated glass bottom cell imaging plates (Eppendorf) and imaged on a DMi8 microscope (Leica Microsystems) with a high-contrast Plan Apochromat 63×/1.30 Glyc CORR CS objective (Leica Microsystems), an ORCA-Flash4.0 digital camera (Hamamatsu Photonics), and MetaMorph 7.8 software (MDS Analytical Technologies). Mitochondrial DNA was labelled with 1 μg/mL DAPI (Sigma-Aldrich) for 20 minutes at 30° C. Cells were washed prior to imaging. For NVJ localization, 2 OD_600_/mL of log-phase yeast were incubated at 30° C for 18 hours in 2 mL synthetic complete media with 2% acetate as the carbon source. Yeast were imaged as above except washed cells were maintained in nutrient-depleted synthetic acetate media during imaging.

All measurements were done on raw images; for presentation, images were processed with MetaMorph 7.8 software. Cropping and adjustment of brightness and contrast was performed in Photoshop CC 2017 (Adobe).

Images were analyzed using custom MetaMorph 7.8 journals. Live cells were identified through several iterations of the Count Nuclei function. Puncta number and size were identified with the Granularity function. Puncta attributed to dead cells and extracellular noise were masked by using the LogicalAND function. The area of colocalization of Vps13^GFP and DAPI-labelled mitochondria, identified with the Granularity function, were determined by using the built-in Measure Colocalization application. The colocalizing Vps13^GFP and NVJ1-RFP structures were determined by masking NVJs identified by the TopHat function in both channels and counted using the Granularity function.

### CPY secretion

To examine the secretion of CPY, 1 OD_600_ of log-phase yeast were serially diluted and spotted on minimal synthetic dextrose media plates. The plates were covered with nitrocellulose and incubated overnight at 30° C. The nitrocellulose membranes were washed to remove cells and blotted with monoclonal mouse anti-carboxypeptidase Y (A-6428; Thermo Fisher Scientific). Blots were developed with West Femto (Pierce) chemiluminescent reagent and exposed to Amersham Hyperfilm (GE Healthcare).

### Sporulation

To examine sporulation efficiency, 0.25 OD_600_ of log-phase diploid yeast were inoculated in 5 mL sporulation media (0.3% KAc, 0.2% raffinose, 0.012% amino acid mix) and incubated at room temperature for 4 days. The cells were washed, diluted and plated on selective media to select for spores. Plates were incubated at 30° C and colonies counted after 3 days.

### Statistical Analysis

Statistical analyses were carried out by performing one-way ANOVA with a Dunnett-corrected post hoc test for comparisons to WT using Prism 6 (GraphPad Software). Data distribution was assumed to be normal, but this was not formally tested.

## Acknowledgements

We thank Dr. Benoît Kornmann (ETH Zürich, Zürich, Switzerland) and Dr. Mike Henne (University of Texas Southwestern Medical School, Dallas, TX) for strains and plasmids and Dr. Luc Berthiaume (University of Alberta, Edmonton, Canada) for his generous gift of rabbit anti-GFP serum.

---

**Supplemental Figure 1:**
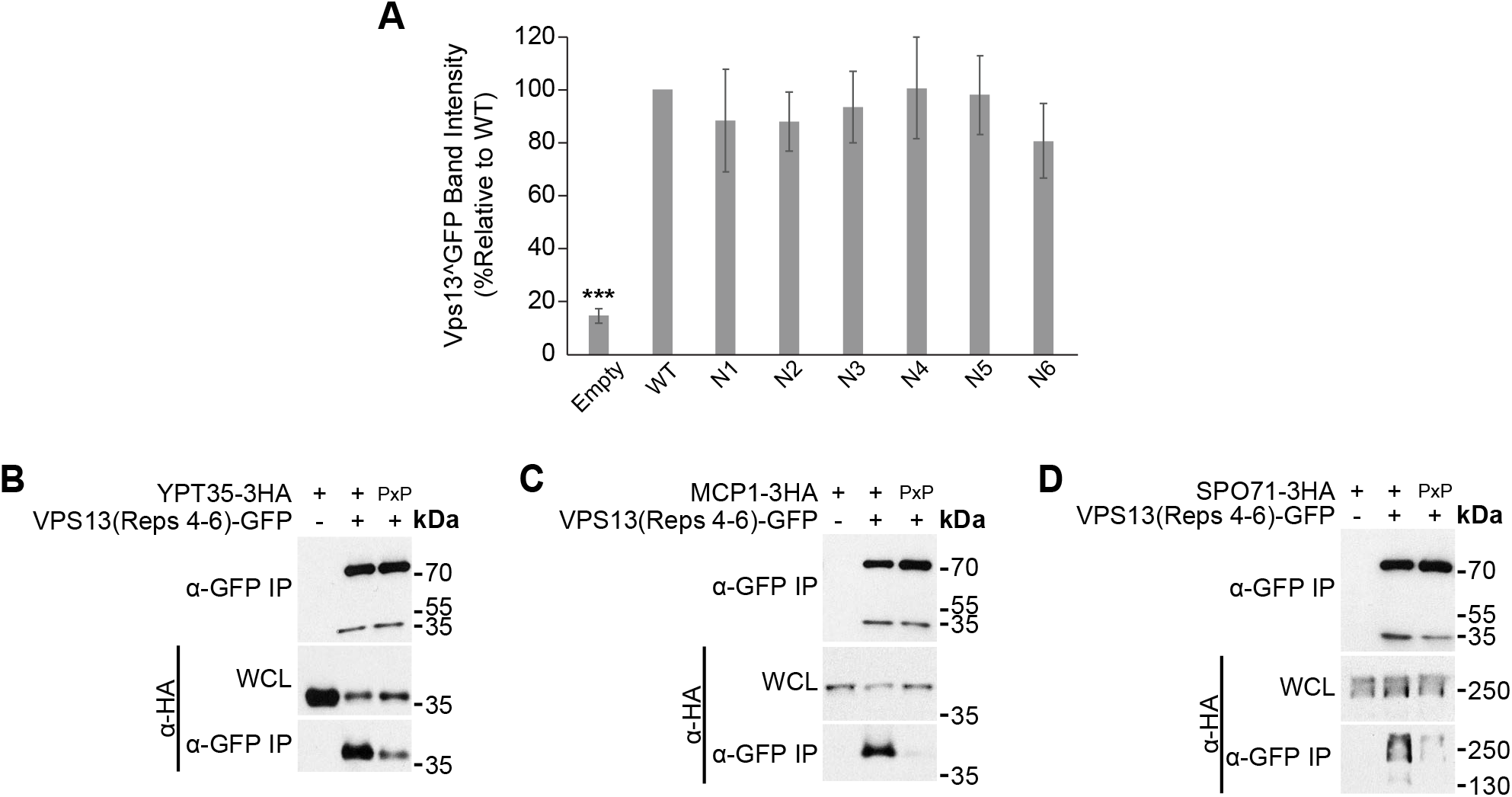
Vps13 VAB mutant stability and Vps13 adaptor binding to VAB repeats 4-6. (A) Recovery of WT and mutant Vps13^GFP from anti-GFP immunoprecipitations analyzed by gel densitometry. One-way ANOVA with Dunneti-corrected post-hoc test; n=11; ***, P <0.001. (B-D) Coimmunoprecipitations of overexpressed adaptors with WT or mutant P×P motifs, as indicated, and *ADH1pr*-driven ENVY-tagged Vps13 VAB repeats 4-6. Proteins were expressed on plasmids in *vps13Δ* strains. IP, immunoprecipitation; WCL, whole-cell lysate. Representative images shown; n = 3.

**Supplemental Figure 2:**
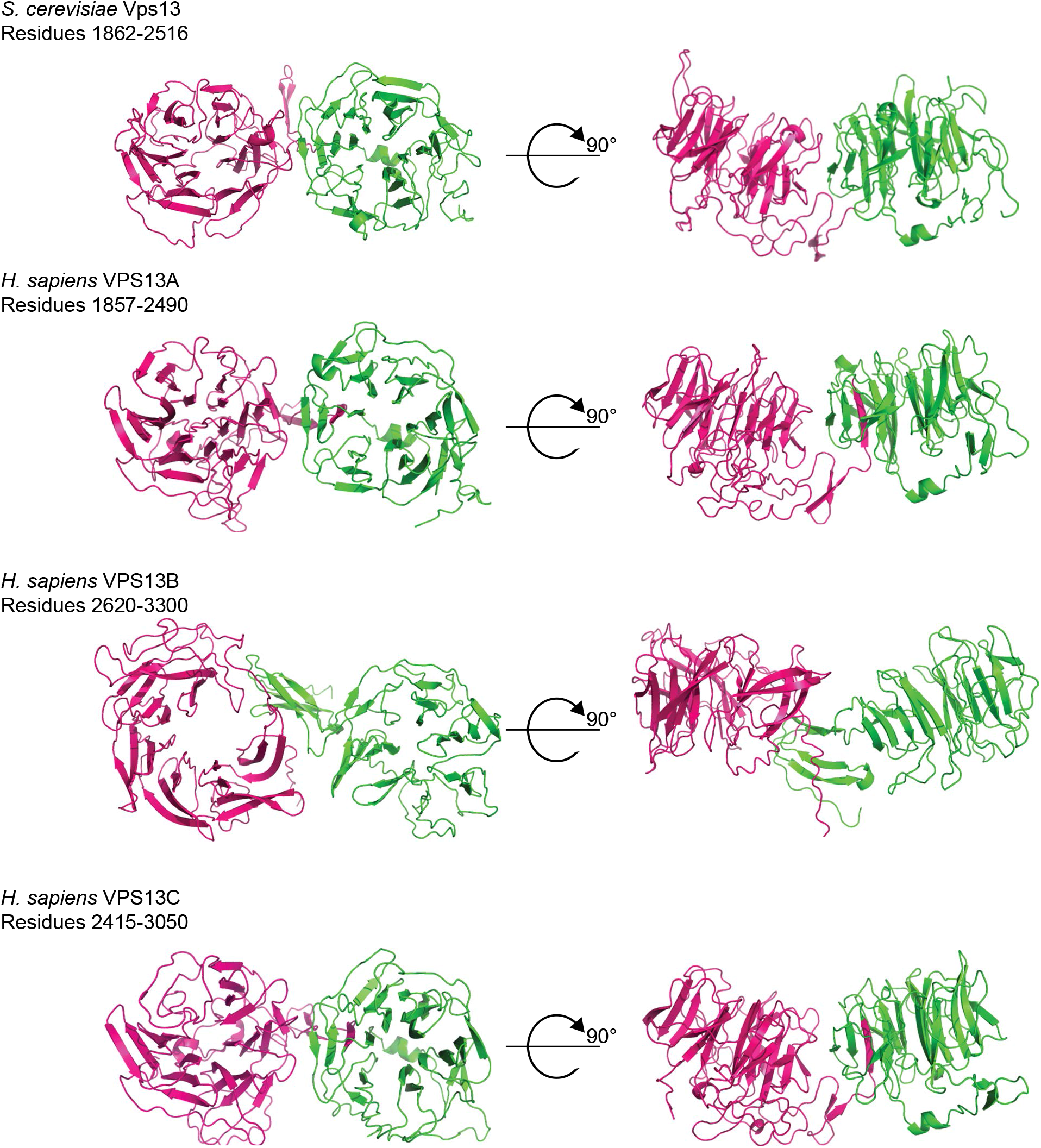
Predicted structural models of yeast and human VAB domains. Homology models of the yeast and human VPS13 VAB domains from I-TASSER (zhanglab.ccmb.med.umich.edu) with modeled residues indicated. VAB repeats 1-3 are shown in pink and repeats 4-6 are shown in green.

